# Circulating ACE2-expressing Exosomes Block SARS-CoV-2 Infection as an Innate Antiviral Mechanism

**DOI:** 10.1101/2020.12.03.407031

**Authors:** Lamiaa El-Shennawy, Andrew D. Hoffmann, Nurmaa K. Dashzeveg, Paul J. Mehl, Zihao Yu, Valerie L. Tokars, Vlad Nicolaescu, Carolina Ostiguin, Yuzhi Jia, Lin Li, Kevin Furlong, Chengsheng Mao, Jan Wysocki, Daniel Batlle, Thomas J. Hope, Yang Shen, Yuan Luo, Young Chae, Hui Zhang, Suchitra Swaminathan, Glenn C. Randall, Alexis R Demonbreun, Michael G Ison, Deyu Fang, Huiping Liu

## Abstract

The severe acute respiratory syndrome coronavirus 2 (SARS-CoV-2) causes the coronavirus disease 2019 (COVID-19) with innate and adaptive immune response triggered in such patients by viral antigens. Both convalescent plasma and engineered high affinity human monoclonal antibodies have shown therapeutic potential to treat COVID-19. Whether additional antiviral soluble factors exist in peripheral blood remain understudied. Herein, we detected circulating exosomes that express the SARS-CoV-2 viral entry receptor angiotensin-converting enzyme 2 (ACE2) in plasma of both healthy donors and convalescent COVID-19 patients. We demonstrated that exosomal ACE2 competes with cellular ACE2 for neutralization of SARS-CoV-2 infection. ACE2-expressing (ACE2^+^) exosomes blocked the binding of the viral spike (S) protein RBD to ACE2^+^ cells in a dose dependent manner, which was 400- to 700-fold more potent than that of vesicle-free recombinant human ACE2 extracellular domain protein (rhACE2). As a consequence, exosomal ACE2 prevented SARS-CoV-2 pseudotype virus tethering and infection of human host cells at a 50-150 fold higher efficacy than rhACE2. A similar antiviral activity of exosomal ACE2 was further demonstrated to block wild-type live SARS-CoV-2 infection. Of note, depletion of ACE2^+^ exosomes from COVID-19 patient plasma impaired the ability to block SARS-CoV-2 RBD binding to host cells. Our data demonstrate that ACE2^+^ exosomes can serve as a decoy therapeutic and a possible innate antiviral mechanism to block SARS-CoV-2 infection.

## Main Text

COVID-19 ^1,2^ has become a global pandemic resulting in more than 55 million cases and over 1.3 million deaths to date ^3^. SARS-CoV-2 shares high structural similarity with SARS-CoV, which caused an outbreak in 2003 ^4^, and encodes four main proteins-glycoproteins spike (S), envelope (E), membrane (M), and nucleocapsid (N) protein besides several accessory proteins ^1,5–7^. Through the external receptor-binding domain (RBD) of the S transmembrane protein, both coronaviruses bind to angiotensin-converting enzyme 2 (ACE2) as a primary receptor for host cell attachment and subsequent entry ^1,5–7^. Approaches to block or impede the viral interaction with the entry receptor ACE2 on the host cell, including S-specific neutralization antibodies (Abs) ^8–20^ and rhACE2 ^21–23^, inhibit infectivity and prevent COVID-19.

Exosomes are cell-secreted extracellular vesicles (EVs) that participate in a variety of physiological and pathobiological functions ^24–28^, and present many proteins on the surface reminiscent of their cellular counterpart, such as immune regulators of myeloid and lymphoid cells to affect antiviral immune response ^28–30^. We detected circulating ACE2^+^ exosomes in plasma from both healthy donors and patients who recovered from COVID-19. Importantly, ACE2^+^ exosomes inhibit SARS-CoV-2 infection by blocking the viral spike protein binding with its cellular receptor in host cells. Our observations demonstrate that ACE2^+^ exosomes are a previously unknown innate antiviral mechanism to prevent SARS-CoV-2 infection, as well as provide a rationale for the use of ACE2^+^ exosomes to combat COVID-19.

### Identification and characterization of ACE2^+^ exosomes

We previously established an automated and high throughput method, micro flow vesiclometry (MFV), to detect and profile the surface proteins of blood exosomes at single particle resolution ^31^. To establish a working protocol for measuring ACE2 expression in exosomes from human plasma, we first determined that ACE2 was detectable in ACE2^+^ cell-derived exosomes. Two sets of human cell lines HEK-293 (HEK) and HeLa, originally negative for ACE2 (ACE2^−^ control), were engineered to stably express ACE2 as verified by flow cytometry and immunoblotting (**Fig. 1A & B**). Both ACE2^+^ cell- and ACE2^−^ cell-derived exosomes were purified from cell culture supernatants using ultracentrifugation after removal of cell debris and apoptotic bodies (**Supplementary Fig. S1A**). Exosomes exhibited an average size of approximately 180 nm with equivalent vesicle counts of 6×10^7^ per μg of exosome proteins as measured by nanoparticle tracking analysis (NTA) (**Fig. 1C**). Immunoblotting demonstrated that these exosomes expressed CD63, CD81, and TSG101 but not GRP94 **(Fig. 1B**).

**Fig. 1.**
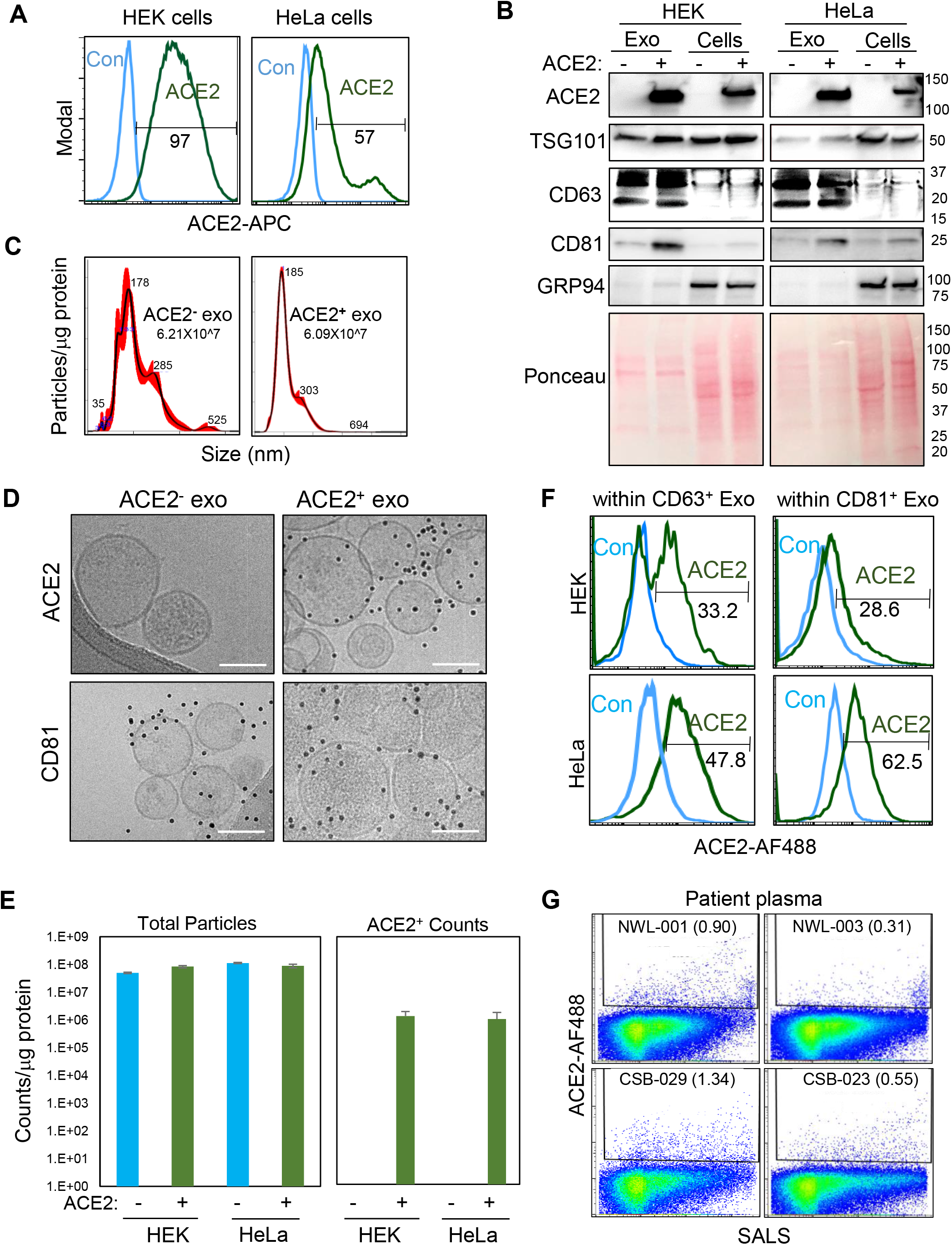
Characterization of ACE2^+^ exosomes. **A**. Flow profiles of ACE2 expression in HEK and HeLa parental control cells (con, light blue line, ACE2^−^) and with ACE2 overexpression (ACE2, green line). **B**. Immunoblots of HEK and HeLa (ACE2^−^ and ACE2^+^) exosomes and cell lysates for ACE2, TSG101, CD63, CD81, GRP94 and loading control of the membrane proteins upon Ponceau staining. **C**. Nanosight-based NTA analysis of the sizes of HEK-derived ACE2^−^ and ACE2^+^ exosomes. **D**. Cryo-EM images of HEK-derived ACE2^−^ (left) and ACE2^+^ (right) exosomes stained with ACE2 (top) and CD81 (bottom panels). Scale bars=100 nm. **E**. Quantified counts of Apogee MFV-based total extracellular vesicles (EV) and ACE2^+^ exosomes. **F**. Overlay flow profiles of ACE2 positivity within CD81^+^ (left panels) and CD63^+^ (right panels) exosomes isolated from HEK-ACE2 (top panels) and HeLa-ACE2 (bottom panels) cells, respectively. **G**. MFV detection of circulating ACE2^+^ exosomes in human plasma of pre-COVID-19 (NWL-001 and - 03) and COVID-19 (CSB-029 and -023).

We further developed two cutting-edge platforms: high-resolution cryogenic electron microscopy (cryo-EM) and high-throughput MFV to analyze native exosomes in liquid conditions at single-exosome resolution. Immuno-cryo-EM revealed that both ACE2^−^ and ACE2^+^ exosomes have a similar spheric vesicle shape and express CD81 (**Fig. 1D**). More importantly, ACE2 expression was detected in a majority of the imaged vesicles (~52% of 178, between 3-40 gold nanoparticles per vesicle) isolated from ACE2^+^ cells **(Fig. 1D, Supplementary Fig S1C & D**). MFV also exclusively detected ACE2 in the exosomes derived from both HEK and HeLa cells expressing ACE2, but not from their parental ACE2^−^ controls, whereas both ACE2^+^ and ACE2^−^ cells produced almost equivalent numbers of total EVs (0.5~1×10^8^ counts per μg exosome proteins), consistent with the NTA analyses (**Fig. 1C & E, Supplementary Fig S1E-F**). Within both ACE2^+^ HEK and HeLa cell-derived exosomes, the ACE2^+^ exosomes account for up to 1-5×10^6^ particles per μg of exosome proteins (2-6% of total EVs) (**Fig. 1E, Supplementary Fig S1F**). Exosomal ACE2 might be under-detected by MFV due to its relatively higher detection threshold than that of cryo-EM images (~50% positivity of ACE2) which provide a resolution of single gold nanoparticles stained for ACE2 (**Supplementary Fig S1D**). The MFV analyses of double stained EVs isolated from HEK-ACE2 and HeLa-ACE2 cells further identified ACE2 largely enriched in CD81^+^ exosomes (28.6 and 62.5%) or CD63^+^ exosomes (33.2 and 47.8%) (**Fig. 1F, Supplementary Fig S1F**).

Taking advantage of the MFV-based direct analysis of circulating exosomes in crude plasma samples, we detected a small subset, ranging from 0.3 to 1.3%, of ACE2^+^ exosomes in total plasma particles or EVs, which were also enriched in CD63^+^ exosome subsets from both pre-COVID-19 (NWL-001 and −004) and COVID-19 (CSB-023 and −029) patients (**Fig. 1G, Supplementary Fig S1G).** The ultracentrifugation pellets of enriched exosomes from plasma specimens had detectable TSG101 and very low ACE2 levels (**Supplementary Fig. S1H**) further confirming that ACE2^+^ exosomes exist in plasma from both healthy donors and COVID-19 patients.

### ACE2^+^ exosomes block SARS-CoV-2 RBD binding and viral infections

To analyze the effects of ACE2^+^ exosomes on viral infection, we implemented a flow cytometry-based SARS-CoV-2 S protein (RBD)-binding assay (**Supplementary Fig. S2A**). As expected, ACE2^+^ HEK cells displayed a specific and high binding (>90%) with a red fluorophore AF-647-conjugated RBD protein at 16 nM, in contrast to a minimal background level of mock control as well as absent RBD-binding with ACE2^−^ control cells (**Supplementary Fig. S2B**). The RBD probe bound to ACE2^+^ cells in a dose-dependent manner, which was inhibited by pre-incubation with rhACE2 as reported previously ^21^ (**Fig. 2A & B**). Notably, ACE2^+^ exosomes inhibited SARS-CoV-2 RBD binding to ACE2^+^ HEK cells as evidence by the percentage of AF-647^+^ cells and the mean fluorescence intensity (MFI) which were both dramatically reduced (**Fig. 2A & B**). In contrast, ACE2^−^ exosomes had negligible effects (**Fig. 2A & B**), indicating that the exosomes inhibit SARS-CoV-2 RBD recognition with their cellular receptors through ACE2. The approximate IC_50_ for soluble rhACE2 to inhibit RBD binding to host cells is 127 nM (~7.6×10^12^ molecules in 0.1 mL of reaction) (**Fig. 2C**). In order to compare the efficacy between rhACE2 and exosomal ACE2 (exoACE2), we converted the exosome concentrations in exosome particles per measured exosome proteins (~1×10^8^/μg) multiplied with estimated number of ACE2 molecules per exosome into molar concentrations of exoACE2. Based on our immune-cryo-EM data, ~50% exosomes presenting 3-40 ACE2 molecules (**Supplementary Fig. S1C-D**), we speculate an average of 20 ACE2 molecules per exosome due to limited exosomal space ^32^. Based upon a series of exosome dilution-based RBD neutralization assays, the IC_50_ values of exoACE2 in the exosomes derived from ACE2^+^ HEK and Hela cells are 0.18 and 0.33 nM, respectively, which contain 1.0-2.0×10^10^ ACE2 molecules in 0.5-1.0×10^9^ particles (**Fig. 2D-E**). Therefore, exoACE2 possesses 400-700 times better neutralization efficiency than soluble rhACE2 to block SARS-CoV-2 viral RBD binding to human host cells.

**Fig. 2.**
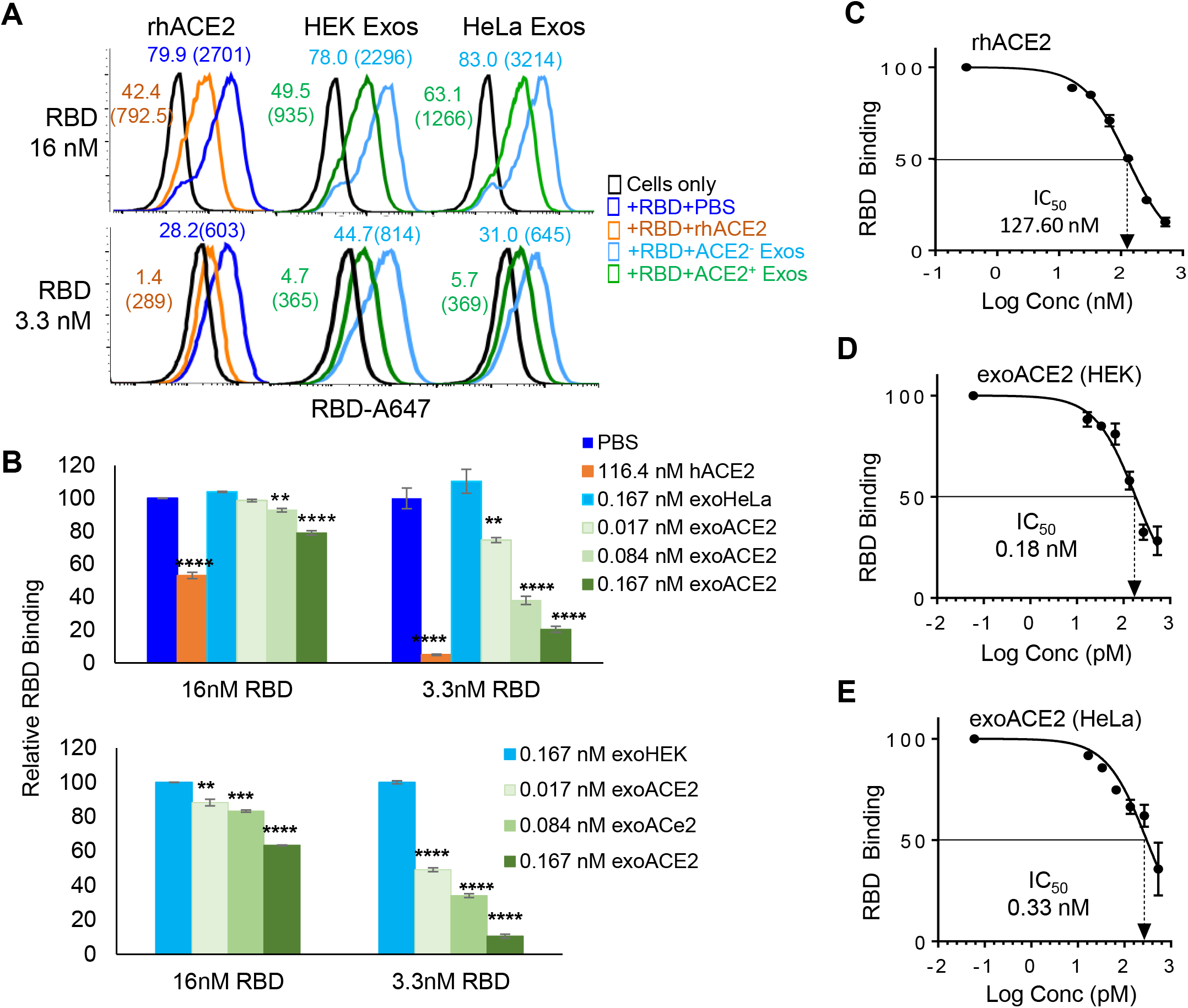
Neutralization effects of ACE2^+^ exosomes on RBD-binding to human host cells. **A**. Flow profiles of RBD binding inhibited by rhACE2 and ACE2^+^ exosomes from HEK-293 and HeLa cells whereas ACE2^−^ exosomes had minimal effects. **B**. Histogram bars of quantified RBD-neutralization by ACE2^+^ exosomes in a dose dependent manner (**p<0.01, ***p<0.001 and ****p<0.0001 compared to PBS or HEK Exos). **C-E.** IC_50_ of rhACE2 and exosomal ACE2 (exoACE2) in the exosomes from ACE2^+^ HEK and HeLa cells on 16 nM RBD-host cell binding (%).

Next, we evaluated the neutralization effects of ACE2^+^ exosomes on the infectivity by SARS-CoV-2 S^+^ pseudovirus and wild-type virus. When the SIV3-derived SARS-CoV-2 S^+^ pseudovirus with either a dual Luc2-IRES-Cherry reporter or an eGFP fluorescence protein reporter was utilized, the viral infectivity to ACE2^+^ host cells was analyzed by flow cytometry of Cherry or eGFP expressing cells as well as luminescence signal of cellular luciferase activity (**Supplementary Fig. S3A-C**). About 3% of ACE2^+^ cells were detected with Cherry protein expression when 300~500 infection-units (IFU) of SARS-CoV-2 S^+^ pseudoviruses were used, whereas no ACE2^−^ cells were infected with the same viral dose (**Fig. 3A, Supplementary Fig. S3D**). ACE2^+^ exosomes blocked SARS-CoV-2 S^+^ pseudovirus infection in a dose-dependent manner and achieved more than 90% inhibition at a dose of 20 μg (**Fig. 3B**). A similar result was observed when the SARS-CoV-2 S^+^ pseudovirus carrying a luciferase reporter was used (**Fig. 3C-E**). The IC_50_ values of exoACE2 in the exosomes derived from ACE2^+^ HEK and HeLa cells are 12.3 pM and 39.7 pM, respectively (**Fig. 3C & D**), which are equivalent to 0.4-1.0×10^8^ particles (0.37-1.18 μg exosomes) with maximal 0.8-2×10^9^ ACE2 molecules in 0.1 mL reactions. In comparison to an IC_50_ of rhACE2 at 1.88 nM (1×10^11^ molecules in 0.1mL), exosomal ACE2 presents an estimated 50 to 150-fold neutralization efficacy to block SARS-COV-2 infection of human host cells. In contrast, ACE2^−^ control exosomes only had marginal neutralization effects, which were not dose-dependent (**Fig. 3F & G**). Preincubation of SARS-CoV-2 S^+^ pseudovirus with ACE2^+^ exosomes did not yield infection of ACE2-negative cells (**Supplementary Fig. S3D**).

**Fig. 3.**
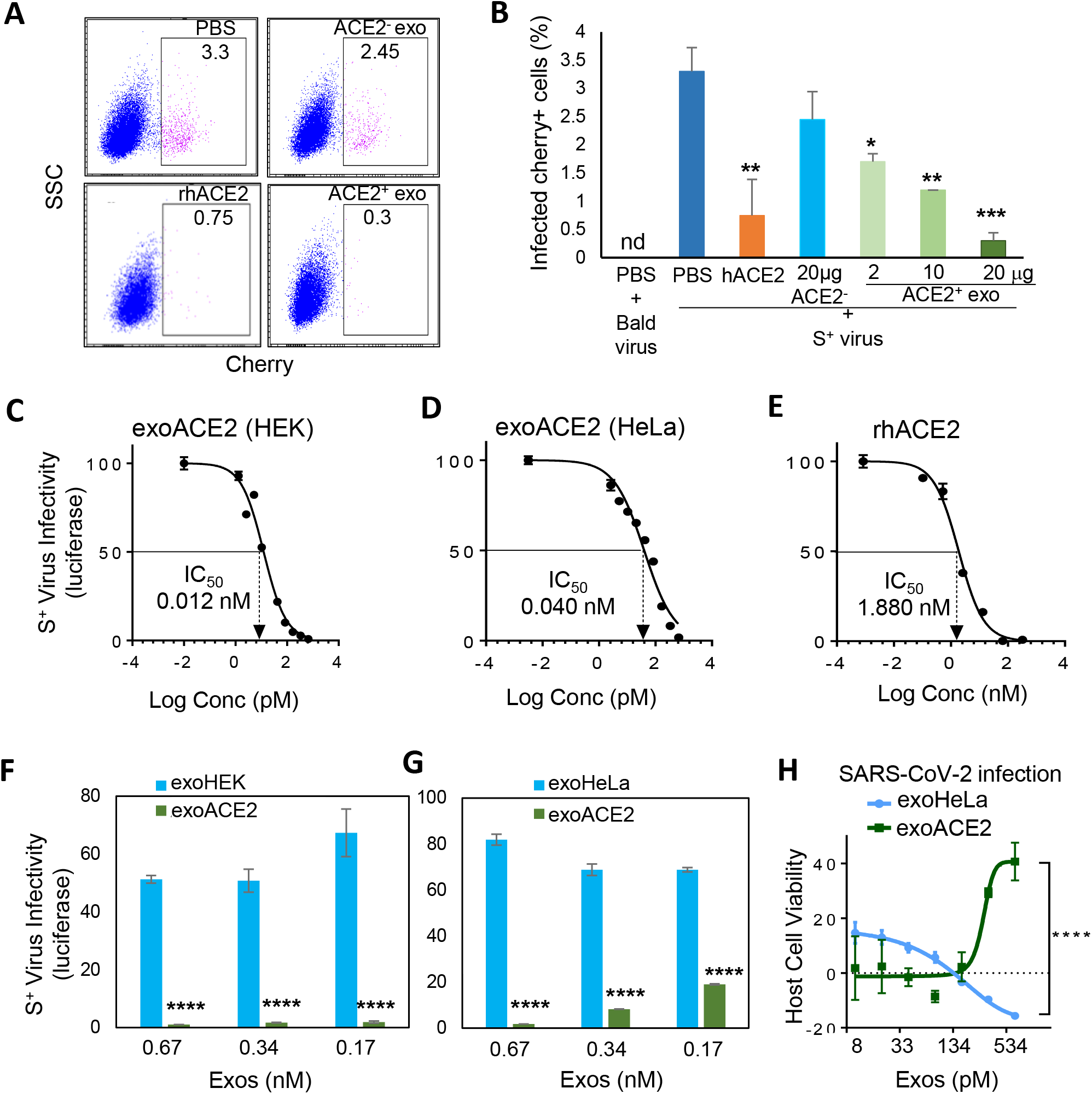
ACE2^+^ exosomes inhibit SARS-CoV-2 S^+^ pseudovirus infection to human host cells. **A-B.** Flow plots (**A**) and bar graph **(B)** of pseudovirus-infected HeLa-ACE2 cells, detected with Cherry reporter expression which was inhibited by rhACE2 and ACE2^+^ exosomes, but not ACE2^−^ exosomes (nd= non-detectable, *p<0.05, **p<0.01 and ***p<0.001 compared to PBS+Spike^+^). **C-E.** IC_50_ of exosomal ACE2 (exoACE2) in ACE2^+^ exosomes from HEK and HeLa cells and rhACE2. **F-G.** Luciferase-based SARS-COV-2 S^+^ pseudotype infectivity of HeLa-ACE2 cells in the presence of ACE2^−^ and ACE2^+^ exosomes from both HEK **(F)** and HeLa **(G)** cells (****p<0.0001 compared to respective ACE2− exosomes). **H.** Vero-6 cell viability upon wild-type SARS-CoV-2 infection which is partially protected by ACE2^+^ HeLa exosomes whereas ACE2^−^ had no effects (****p<0.0001).

Collectively, our results provide a rationale for the use of ACE2^+^ exosomes as an innovative methodology to prevent SARS-CoV-2 infection. Indeed, upon the wild-type SARS-CoV-2 infection (400 plaque-forming units), HeLa ACE2^+^ exosomes at the doses of 334 and 668 pM exoACE2 (10 and 20 μg exosomes) protected the vero-6 cells from viral infection-mediated death resulting in an improved host cell viability whereas ACE2^−^ exosomes failed to protect the cells from viral infection (**Fig 3H**).

### ACE2^+^ exosomes associated with plasma neutralization effects

We continued to investigate whether ACE2^+^ exosome abundancy is associated with viral neutralization effect of human plasma. MFV-based single exosome analysis detected a wide range of ACE2^+^ exosome abundancy (vesicle counts) in plasma from both pre-COVID-19 (NWL) and convalescent COVID-19 (CSB) patients (**Figs. 1G & 4A),** implying that the levels of circulating ACE2^+^ exosomes are variable. As expected, the RBD-IgG levels in COVID-19 patient plasma were significantly associated with their neutralization effects on RBD binding to human cells, but pre-COVID-19 plasma (NWL) had absent or negligible levels of RBD-IgG **(Fig. 4B, Supplementary Fig. S3E)**.

**Fig. 4.**
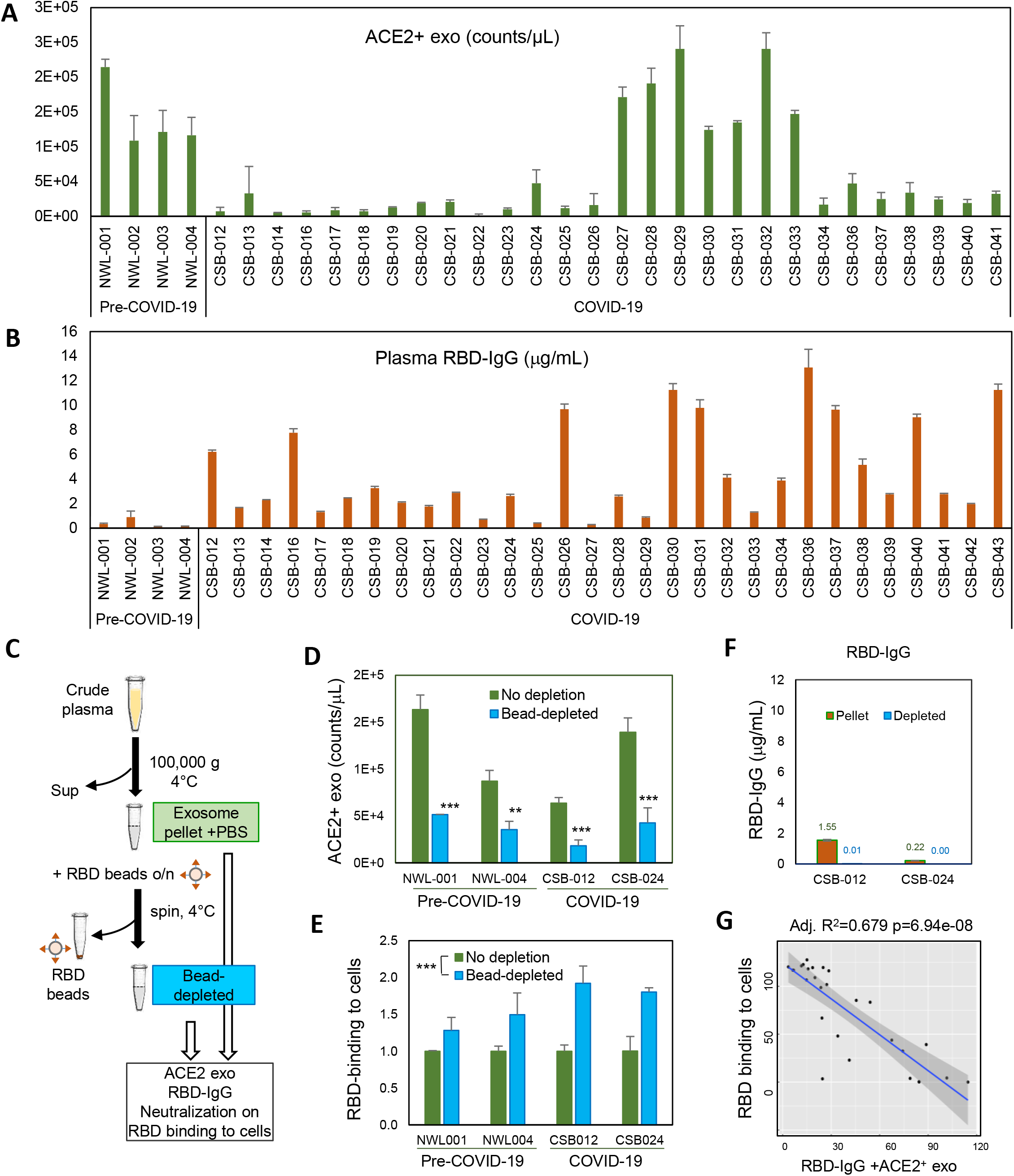
Detection of ACE2^+^ exosomes in NWL and CSB patient plasma. **A-B.** MFV counts of ACE2^+^ exosomes and ELISA-measured RBD-IgG levels in patient plasma, pre-COVID-19 (NWL) and convalescent COVID-19 plasma (CSB) **C.** Schematic of exosome enrichment and RBD-beads based depletion. **D-E.** Levels of ACE2^+^ exosome counts (**D**) and altered RBD-host cell binding **(E)** of the plasma exosome pellet prior to and after RBD-bead depletion (**p<0.01 and ***p<0.001). **F.** RBD-bead depletion reduced the residual RBD-IgG cells. **G.** Negative association of RBD binding to host cells (ACE2^+^ HEK) with the integrated RBD-IgG and ACE2^+^ exosome levels in the convalescent COVID-19 patient plasma (N=30). ACE2^+^ exosomes and RBD-IgG levels were integrated using their coefficients following a linear regression.

Together with our discovery that ACE2^+^ exosome inhibit SARS-CoV-2 infection, we speculated that, in addition to specific neutralization Abs, ACE2^+^ exosomes might function as an innate anti-SARS-COV-2 mechanism. To investigate this idea, we first enriched plasma exosomes by ultracentrifugation (**Fig. 4C & D**). Importantly, ACE2^+^ exosomes pelleted from the plasma of both healthy donors and COVID-19 patients partially inhibited RBD-binding to ACE2-expressing HEK cells (**Fig. 4E**). To validate whether the ACE2^+^ exosomes in the plasma pellet were responsible for inhibition of RBD binding, we used RBD-conjugated magnetic beads to deplete the majority of ACE2^+^ exosomes in the samples (**Fig. 4C & D**). Depletion of ACE2^+^ exosomes significantly impaired the ability of plasma samples to inhibt RBD-binding to ACE2^+^ HEK cells (**Fig. 4E**), indicating that the ACE2^+^ exosomes in the plasma from both healthy donors and COVID-19 patients were responsible for anti-SARS-CoV-2 activity.

We also noticed that the pellets from one COVID-19 plasma (CSB-012) had residual but detectable levels of RBD-specific IgG, which were also depleted by RBD-beads (**Fig. 4C & F**). This implies that IgG may contribute to the neutralization capacity of the plasma pellets from COVID-19 patients, but not the healthy donors, to suppress SARS-CoV-2 infectivity. Based on the altered ACE2^+^ exosome levels (vesicle counts measured by MFV) in all four samples and the RBD-IgG alteration in CSB-012 and CSB-024, we developed a mathematical model to estimate the relative contributions of ACE2^+^ exosomes and RBD-IgG in convalescent plasma to block SARS-CoV-2 binding (**Supplementary Fig. S3F**). We then integrated the contribution of both ACE2^+^ exosomes and RBD-IgG levels for an improved prediction of RBD neutralization activity of COVID-19 patient plasma (N=30) (**Fig. 4G, Supplementary Fig. S4**).

Collectively, our data reveal a previously unknown antiviral mechanism of circulating ACE2^+^ exosomes that may suppress infection by SARS-CoV-2 and other viruses utilizing ACE2 as an entry receptor.

## Discussion

Our studies demonstrate that ACE2^+^ exosomes can compete with host cell surface ACE2 to inhibit SARS-CoV-2 infection, and possibly by other coronaviruses that utilize ACE2 as their initial tethering receptor. Similarly, a recent study has shown that rhACE2 inhibits SARS-CoV-2 infection ^21^. Based on an assumption that one exosome is capable of containing a limited number of total protein molecules ^32^ and our cryo-EM data showing possibly up to 20 ACE2 molecules per exosome, exosomal ACE2 might possess up to 100-fold better efficiency to block SARS-CoV-2 infection than soluble rhACE2.

Exosomes have been utilized as drug delivery systems with therapeutic potential against various disorders including infectious diseases ^33,34^. We anticipate that the therapeutic efficacy of ACE2^+^ exosomes can be further potentiated through co-delivering additional anti-SARS-CoV-2 medicines ^35,36^. In addition, circulating exosomes in plasma represent an important component of blood in terms of its defensive, homeostatic and signal transduction properties ^28–31^. Importantly, here we detected a substantial amount of ACE2^+^ exosomes in human plasma, and ACE2^+^ exosomes function as a potent decoy to protect host cells from SARS-CoV-2 infection. Our discovery reveals that circulating ACE2^+^ exosomes may serve as a previously unknown innate antiviral mechanism to protect the host from coronavirus infection. Future studies are needed to investigate whether circulating ACE2 exosomes account for reduced SARS-CoV-2 viral load and COVID-19 pathogenesis.

## Acknowledgments

We are thankful to the team of Northwestern COVID-19 Antibody and Cancer Collaborative Group (U54 and U18 Prep Team) and advisory members with their scientific input and resourceful support for the project, especially Drs. Alfred L. George Jr. who also edited our manuscript, James Crowe, Judith Varner, Richard D’Aquila, Leonidas C. Platanias, Rex L. Chishom, Alan R. Hauser, Elizabeth M. McNally, and William A. Muller. The work was partially funded by Northwestern University Feinberg School of Medicine Emerging and Re-emerging Pathogens Program (EREPP) (H.L.), Department of Pharmacology Start-up fund (H.L.), and the R.H. Lurie Comprehensive Cancer Center Blood Biobank fund (M.I.). We gratefully acknowledge the support from the R.H. Lurie Comprehensive Cancer Center Structural Biology Facility and Flow Core. The Ametek K3 DDE at Northwestern University for cryo-EM was generously provided by Professor Robert A. Lamb, Ph.D., Sc.D., HHMI investigator. The ACE2-expressing HeLa cells were provided by Dr. Thomas Gallagher of Stritch Medical School, Loyola University. We appreciate the effort of the BSL-3 facility at the NIAID-supported University of Chicago Howard T. Ricketts Regional Biochontainment Laboratory for performing the wild-type SARS-CoV-2 live virus study.

## Methods

### Human subject study and biosafety approvals

All research activities with human blood specimens of pre-COVID-19 and COVID-19 convalescent patients were implemented under NIH guidelines for human subject studies and the protocols approved by the Northwestern University Institutional Review Board (STU00205299 and #STU00212371) as well as the Institutional Biosafety Committee.

### Cell culture

The parent ACE2^−^ human embryonic kidney HEK-293 cells (HEK) or human cervical cancer HeLa cells (HeLa) are transduced with lentiviral pDual-ACE2 expression vector for stable ACE2 expression and production of ACE2^+^ exosomes. Dr. Thomas Gallagher of Stritch Medical School, Loyola University kindly provided HeLa and HeLa-ACE2 cells via the Hope group. ACE2-parent cell serve as negative controls in production of ACE2-exosomes in the culture. Cells were tested for mycoplasma contamination before culturing. Cells were grown in Dulbecco’s Modified Eagle’s Medium (DMEM) supplemented with 10% (v/v) fetal bovine serum (FBS), 100 U/mL penicillin and 100 mg/mL streptomycin. FBS used to prepare complete media was exosome-depleted by ultracentrifugation at 100,000 × *g* for 16 h at 4 °C.

### Flow cytometry

Cells were blocked with mouse serum IgG (Sigma, 15381) for 10 min at room temperature and then incubated with specific antibodies; AF-647 mouse anti-human ACE2 (R&D systems, FAB9332R), AF-488 mouse anti-human ACE2 (R&D systems, FAB9333G), AF-647 isotype control mouse IgG2b (R&D systems, IC003R) or AF-488 isotype control mouse IgG2bAF488 (R&D systems, IC003G) for 45 min on ice, followed by washing twice with 2% exosome-free FBS/PBS. Finally, the cells were diluted in 2% exo-free FBS/PBS and analyzed on a BD-LSR II flow cytometer (BD Biosciences).

### Isolation and Purification of cell line-derived exosomes

Exosomes were isolated from the cell culture supernatant of each of the four cell lines as described previously^37^. Cells were cultured as monolayers for 48-72 h under an atmosphere of 5% CO_2_ at 37 °C. When cells reached confluency of approximately 80-90%, culture supernatant was collected, and exosomes were isolated using differential centrifugation. First, the supernatant was centrifuged at 2,000 × g for 10 min then at 10,000 × g for 30 min to remove dead cells and cell debris. Second, the supernatant was ultracentrifuged for 70 min at 100,000 × g to pellet the exosomes. Exosomes were then washed by resuspension in 30 mL of sterile PBS (Hyclone, Utah, USA), and pelleted by ultracentrifugation for 70 h at 100,000 × g. The exosome pellet was resuspended in 100 μl PBS and stored at −80 °C.

### Western blotting

Cells and exosomes were lysed by RIPA buffer with protease inhibitor cocktail (1:100 dilution) for 30 min on ice, then centrifuged for 15 min at 4 °C and 14,000 rpm. 10-20 μg of cell-derived proteins and 2-8 μg of exosome-derived proteins were denatured at 100 °C for 5 min and loaded to SDS-PAGE, then transferred to PVDF membranes. The antibodies, ACE2 (R&D systems, AF933), CD81 (GeneTex, CTX101766), CD63 (Proteintech, 25682-1-AP), GRP94 (Proteintech, 1H10B7), TSG101 (Proteintech, 14497-1-AP) and β-actin (Sigma, A5441) were used as primary antibodies and horseradish peroxidase (HRP)-conjugated secondary antibodies were from Promega (Rabbit W401B and Mouse W402B), and the substrate ECL was detected by Pierce ECL2 solution (Thermo Fisher Scientific, 1896433A).

### Nanoparticle tracking analysis

Analysis was performed at the Analytical bioNanoTechnology Core Facility of the Simpson Querrey Institute at Northwestern University. All samples were diluted in PBS to a final volume of 1 ml and ideal measurement concentrations were found by pre-testing the ideal particle per frame value. Settings were according to the manufacturer’s software manual (NanoSight NS3000).

### Micro Flow Vesiclometry (MFV) Analysis of Exosomes

Antibody solutions were centrifuged at 14000 × g for 1 h at 4 °C to remove aggregates before use. Exosomes (1-2 μg protein equivalent amount of exosomes in 20 μL of PBS) were blocked using 1μg of mouse serum IgG for 10 min at RT then incubated with: AF-488 mouse anti-human ACE2 (R&D systems, FAB9333G), APC mouse antihuman CD81 (BD Biosciences, 561958), AF-647 mouse antihuman CD63 (BD, Biosciences, 561983), AF-488 isotype control mouse IgG2b (R&D systems, IC003G), APC isotype control mouse IgG_1_κ (BD Biosciences, 555751) or AF-647 isotype control mouse IgG_1_κ (BD, Biosciences, 557714) for 45 min at 4°C. The solution was then diluted to 200μL with PBS and the samples were run on Apogee A50 Micro Flow Cytometer (MFC) (Apogee Flow Systems, Hertfordshire, UK) (http://www.apogeeflow.com/products.php). The reference ApogeeMix beads (Apogee Flow Systems, 1493), were used to assess the performance of Apogee MFC, and to compare the size distribution of the exosomes. PBS was run as a background control.

### Immuno-cryo-EM imaging

Antibody solutions and other staining buffers were centrifuged to remove non-specific particles or aggregates in the buffer of intereat, at 14000 × g for 1 h at 4 °C before use. Exosomes (10μg in 100uL) were blocked using 5μg of mouse serum IgG for 10 min at RT then incubated with mouse anti-human ACE2 (R&D systems, FAB9333G), mouse antihuman CD81 (BD Biosciences, 551108), isotype control mouse IgG2b (R&D systems, IC003G) or isotype control mouse IgG_1_κ (BD Biosciences, 551954) for 45 min at 4 °C.. To rinse samples, 1mL PBS was added to the tubes, and exosomes were centrifuged 100,000 × g for 30 min at 4°C. PBS was aspirated, samples were reconstituted in 100 uL PBS, and incubated with EM goat anti-mouse IgG (H&L) 10 nm gold conjugated (BBI solutions, EM.GMHL10) (7:100) for 30 min at RT. Exosomes were then rinsed by adding 1300 uL PBS then centrifugation 100,000 × g for 15 min at 4°C. Finally, PBS was aspirated, and exosomes were reconstituted in 50 uL PBS.

For cryoEM visualization, samples were prepared from freshly stained exosomes at the concentration provided. For cryo-freezing, 3.5μl of exosome solutions were applied to fresh glow-discharged (10 s, 15 mA; Pelco EasiGlow) lacey carbon TEM grids (Electron Microscopy Services) and vitrified using a FEI Vitrobot Mark IV (FEI, Hillsboro, OR). The sample was applied to the grid and kept at 85% humidity and 10 oC. After a 10 s incubation period the grid was blotted with Whatman 595 filter paper for 4 seconds using a blot force of 5 and plunge frozen into liquid ethane. Samples were imaged using a JEOL 3200FS electron microscope equipped with an omega energy filter operated at 200 kV with a K3 direct electron detector (Ametek) using the minimal dose system. The total dose for each movie was ~20 e-/A2 and was fractionated into 14 frames at a nominal magnification between 8,000 to 15,000 (pixel size on the detector between 4.1 Å to 2.2 Å, respectively). After motion correction of the movies ^38^, exosomes were identified manually using ImageJ ^39^. Two grids were prepared and imaged with 10-20 fields for each condition.

### Development of the SARS-Cov-2 RBD “bait”

RBD of 223 amino acid (Arg319-Phe541) fragment of the SARS-CoV-2 Spike protein that binds to the ACE2 receptor (Raybiotech, 230-30162-100) was biotinylated using NHS-PEG4-Biotin (Thermo Fisher, 21330). The protein was de-salted using Zeba Quick Spin columns (Thermo Fisher, 89849) and incubated with Streptavidin-AlexaFluor-647 (SA-AF-647) (Thermo Fisher, S21374) to make the RBD-biotin-AF647 bait.

### Cell-based RBD binding neutralization by ACE2^+^ exosomes and human plasma

The RBD-biotin-AF647 bait (3.3 and 16 nM) was incubated with exosomes (ACE2^+^ and ACE2^−^), recombinant human ACE2 extracellular region (rhACE2, RayBiotech, 230-30165), or human plasma (10 μl or 80 μl) for 45 minutes on ice (creating “neutralized RBD”), then incubated with ACE2^+^ HEK-293 cells (200,000 cells in 100 μL) for 45 minutes on ice. Human recombinant ACE2 protein was used as a positive control (70-140 ng). RBD bait that was incubated with PBS, or with ACE2^−^ exosomes, non-fluorescent RBD bait (mock control) and ACE2^−^ cells were used as controls. Cells were then spun and washed twice with PBS. DAPI was added as to exclude dead cells analyzed on flow cytometer and viable singlets were gated for percentage and mean fluorescence intensity (MFI) measurements of the RBD-AF647^+^ population.

### Neutralization effects of ACE2^+^ exosomes on SARS-CoV-2 spike^+^ pseudovirus infection to human host cells

The SARS-CoV-2 spike (S^+^) pseudovirus carrying the Luc2-Cherry reporters were made for live virus neutralization assay after the pcDNA3-spike expression vector was transfected along with pCMV-Luc2-IRES-Cherry and pSIV3+ lentiviral vectors into a lentivirus producing cell HEK-293. The S^+^ pseudovirus was then incubated with ACE2^+^ exosomes, or ACE2^−^ exosomes, or a positive control rhACE2, or negative control (PBS), for 1 hr at 37°C prior to the infection with ACE2^+^ human host cells HeLa in 96-well plates (5,000 cells/well). A bald virus without spike expression and ACE^−^ cells served as negative controls. Flow cytometry of Cherry or eGFP and luciferase activity analysis (Promega, E1500) were used to assess viral infectivity.

### Wild-type SARS-CoV-2 live virus infection to vero-6 cells at the BSL3 facility

The wild-type SARS-CoV-2 live virus study was conducted at the BSL-3 facility at the NIAID-supported University of Chicago Howard T. Ricketts Regional Biochontainment Laboratory. One day prior to viral infections, 10,000 vero-6 cells were seeded per well in triplicates onto 96-well plates. 16 hours after seeding, the attached cells were infected with mock controls (no virus) and wild-type SARS-CoV-2 (500 pfu) viruses which were pre-mixed with a serial of doses of exosomes (starting from 20 μg with 6 times of 1:2 dilutions) or an untreated control. 96 hours later, the host cell viability (opposite to viral infectivity-caused cell death) was measured by crystal violet staining which stained attached viable cells on the plate following fixation. Cells killed off by the virus were floating and excluded. For the untreated control, the cells were infected but left without any treatment with a value of maximal cell death caused by the virus. The second control was the mock infected control where cells grew in the absences of virus or experimental sample representing the maximum normal cell growth over the time period. The absorbance value of the untreated control was subtracted from all other absorbance values thereby making untreated 0 then all absorbance values were divided by the mock infected value thereby making that value 100.

### RBD-IgG quantitative ELISA assay

The ELISA protocol was validated as previously described ^40,41^ and used herein with the modification of using plasma instead of serum. Plasma samples were diluted by half with PBS during RosetteSep human B cell processing (StemCell Technologies #15064), aliquoted, and stored at −80C until analysis. Plasma was run in quadruplicate and reported as the average. Results were normalized to the CR3022 antibody with known affinity to RBD of SARS-CoV-2 ^42^. Sample anti-RBD IgG concentration reported as μg/ml was calculated from the 4PL regression of the CR3022 calibration curve. A sample value >0.39 μg/ml CR3022 was considered seropositive.

### Exosome enrichment by ultracentrifuge

Covid-19 (CSB) and pre-COVID-19 (NWL) patient derived plasma samples were obtained from Northwestern Memorial Hospital and stored at −80 °C. CSB and NWL plasmas were ultra-centrifuged (Beckman Coulter) at 100000 x g for 2, 4, 8 and 18 h at 4 °C to isolate and enrich exosomes in the pellets. After centrifuge, supernatants and plasma pellets were collected separately. Plasma pellets were resuspended in appropriate volumes of PBS. The levels of ACE2^+^ exosomes in plasma samples were evaluated by MFV on Apogee and western blotting using exosome marker TSG101 and ACE2. ACE2^+^ cell line derived exosomes were used as a positive control.

### Depletion of ACE2^+^ exosomes by RBD-beads

CSB and NWL patient plasma samples (1.0-2.0 ml) were ultra-centrifuged for 8 h at 100000 × g at 4 °C, and the pellets were resuspended in 250-500 μL PBS as exosome-enriched samples for subsequent bead-mediated depletion. RBD-coupled magnetic beads or anti ACE2-coupled dynabeads beads were prepared according to manufacturer’s protocols. Biotin conjugated RBD protein (ACROBiosystems, SPD-C82E9) were coupled with magnetic beads (CELLection Biotin Binder Kit, Thermo Fisher Sceintific, 11533D), and 5 μg of anti-ACE2 antibody (R&D systems, AF933) or isotope control IgG (R&D systems) were conjugated to dynabeads (Dynabeads Co-Immunoprecipitation Kit, Thermo Fisher Scientific, 14321D). Exosome pellet samples were incubated with the beads for overnight at 4°C and then the beads were removed by spinning or magnetic forces. The ACE2^+^ exosome depletion efficiency was confirmed by MFV on Apogee and/or western blotting methods.

The altered neutralization effects of NWL and CSB plasma-enriched exosomes (resuspended pellets) prior to and after bead depletion were measured via flow cytometry as modified RBD binding to human host cells as described above. And rhACE2 protein (RayBiotech, 230-30165) was used as a positive control (70-140 ng).

### Statistical Analysis

GraphPad Prism 6.0 Software was used to perform statistical analyses and calculate the IC_50_. One way or two ways ANOVA (followed by Tukey or Sidak’s posttest) were used where appropriate. Data are presented as mean ± standard deviation (SD). For patient plasma association analyses, the neutralization data were collected from the laboratory, electronically recorded and verified by laboratory staff. To reduce bias from batch effects, four different RBD-IgG replicates were performed. One-way ANOVA test was performed showing the replicates did not show significant batch errors (F=0.01, p-value>0.9), thus mean values were used for analysis. Log-linear model (Poisson regression) was fitted to estimate the associations between counts of RBD binding to cells and independent predictors of interest. This suggested negative associations and the adjusted R^2^ suggests that integrated ACE2+RBD value explains the relation better than RBD alone (Adj. R^2^ =0.623 p<0.0001). Coeffecient from this analysis was used to create graphs in Fig 4g. All statistical analyses were performed by R 4.0.2.

---

**Supplementary Fig. S1.**
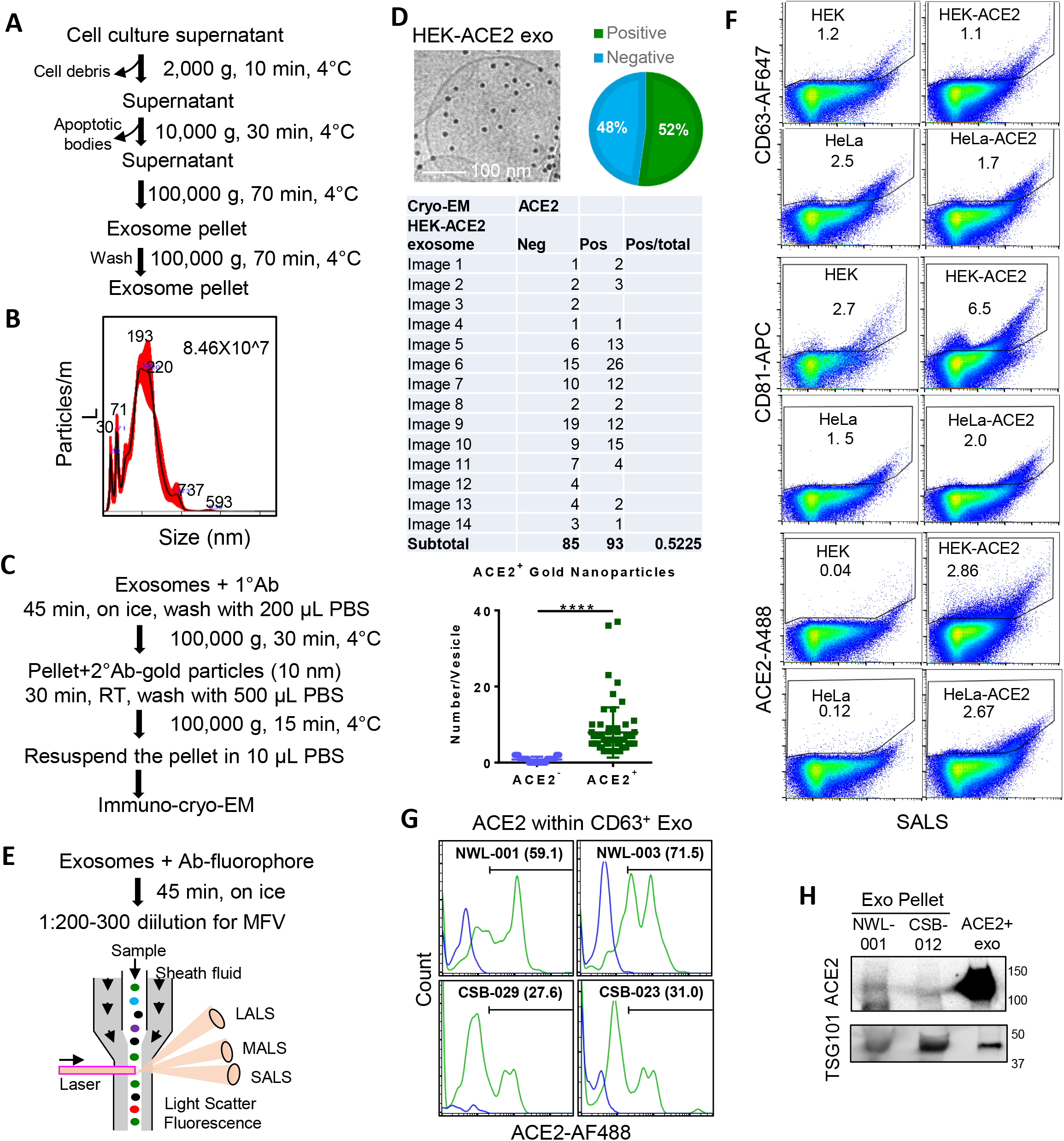
Characterization of ACE2^+^ exosomes. **A.** Exosome isolation protocol and analysis. **B.** Nanosight-based NTA analysis of HeLa-ACE2 cell derived exosomes. **C.** Immuno-cyro-EM staining protocol. **D.** Cryo-EM images of ACE2^+^ exosomes and a table/pie graph of counted HEK-ACE2^+^ and ACE2^−^ exosomes (****p<0.0001). **E.** Staining protocol for Apogee-based MFV on fluorescent and light scatter panels, SALS, MALS, and LALS. **F.** MFV profiles of exosomal CD63, CD81 and ACE2 expression in four cell line-derived exosomes (HEK and HeLa with or without ACE2 expression). **G.** Flow profiles of ACE2 positivity within gated CD63^+^ exosomes directly detected in human plasma (NWL-001, -003 and CSB-029, -023). **H.** Immunoblots of ACE2 and TSG101 with the exosome pellets ultracentrifuged from human plasma (100,000 g × 8 h), NWL-001 and CSB-012.

**Supplementary Fig. S2:**
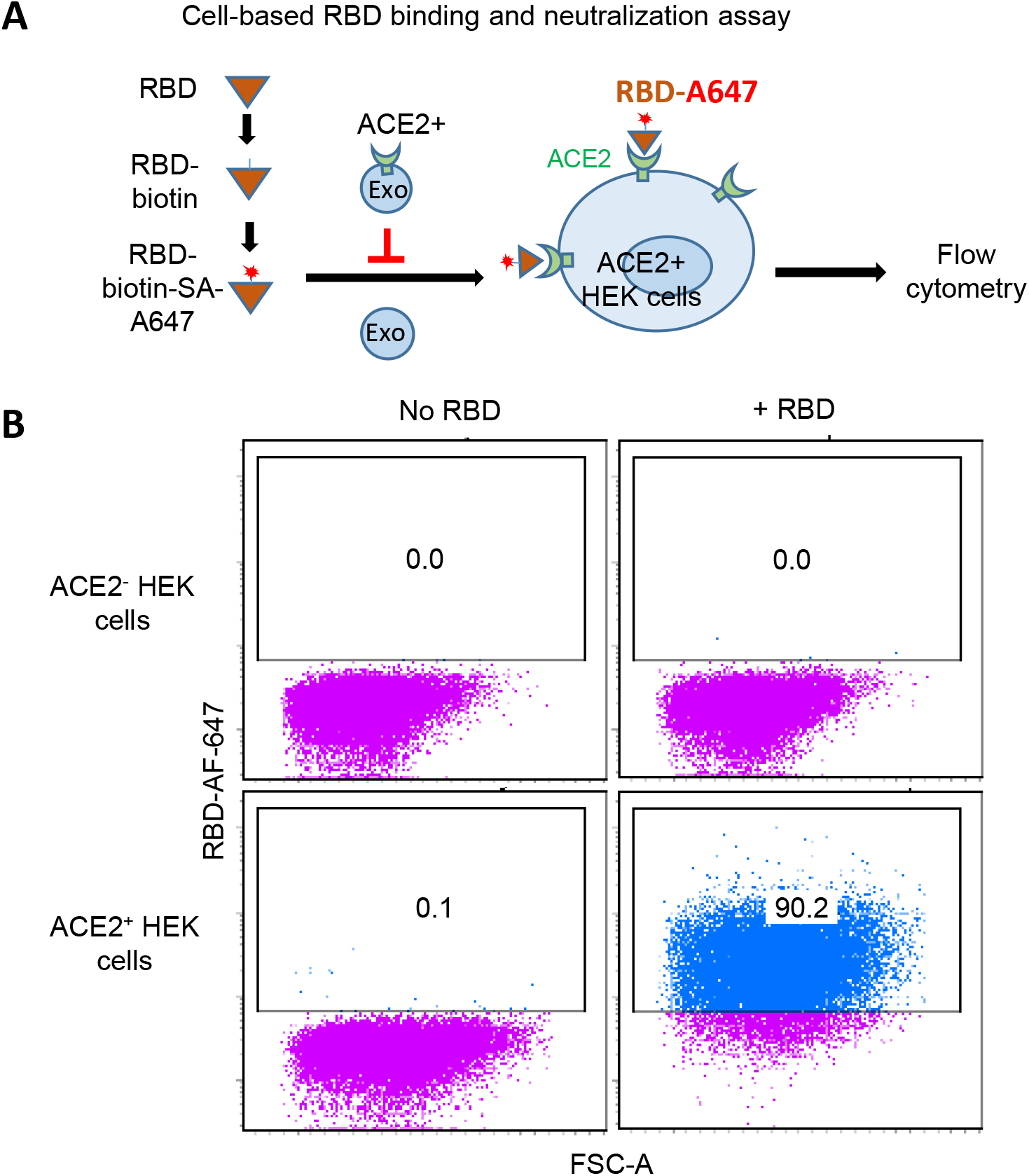
ACE2^+^ exosomes block RBD binding with host cells. **A.** Schematic of flow cytometry-based RBD binding to human host cells (ACE2^+^ HEK) for neutralization effect analysis **B.** Flow plots of ACE2^+^ and ACE2^−^ HEK cells in the absence and presence of AF647-conjugated RBD binding with minimal binding to a mock control of the RBD probe.

**Supplementary Fig S3:**
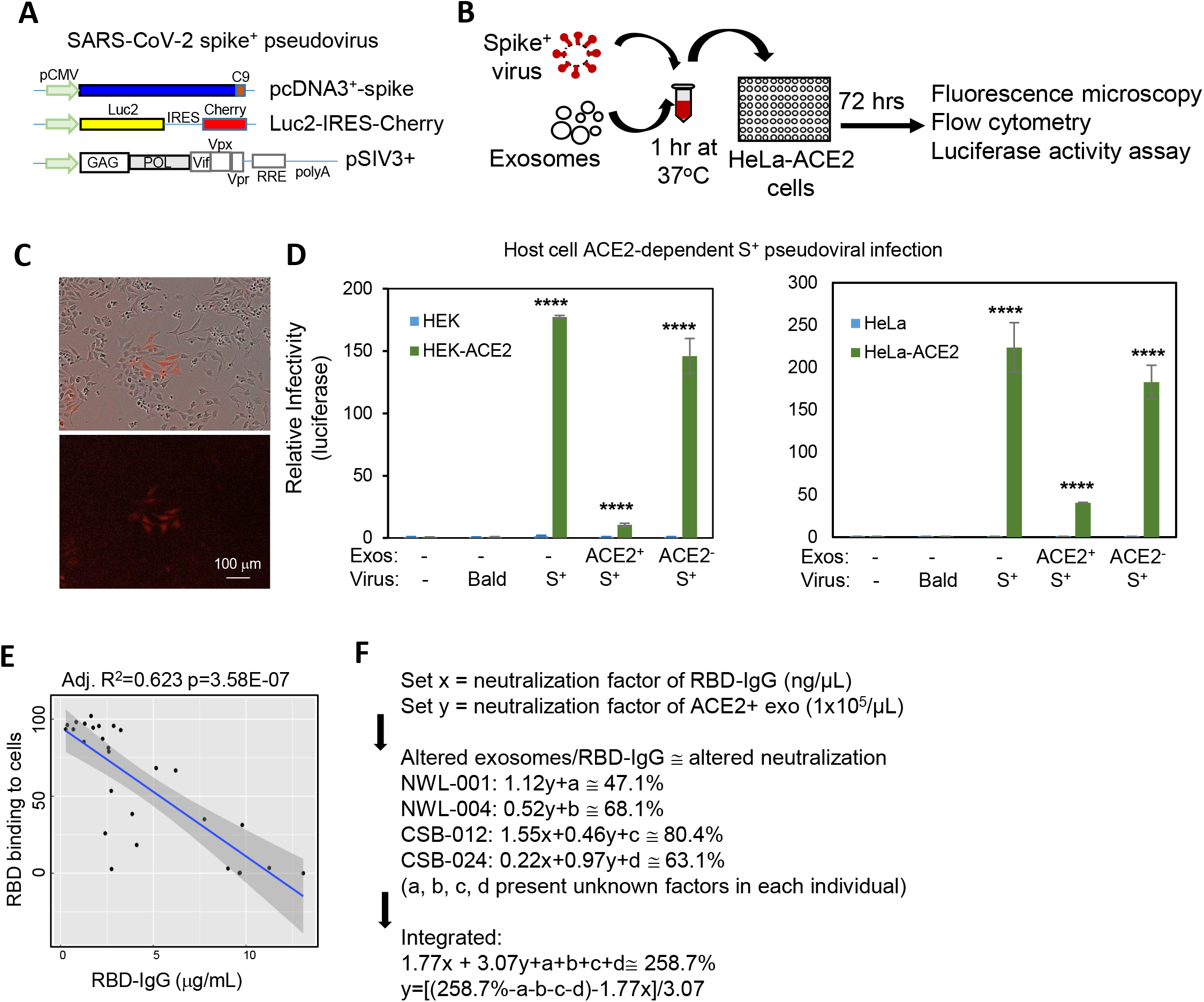
ACE2^+^ exosomes block pseudotyped SARS-CoV-2 infections. **A.** Schematic of the three vectors for SARS-CoV-2 S^+^ pseudovirus production. **B**. Pseudovirus infection with ACE2^+^ HeLa cells and subsequent infectivity analyses, including Cherry positive cells under fluorescent microscopy, MFV, and luciferase activity assays. **C**. Fluorescent images of Cherry^+^ red cells as pseudovirus-infected cells. **D**. Luciferase-based viral infectivity between HEK control and HEK-ACE2 cells (left panel) and between HeLa control and HeLa-ACE2 cells (right panel), 48-72 hr after incubation with bald (S^−^) or S^+^ pseudotyped viruses in the absence and presence of ACE2^+^ or ACE2^−^ exosomes (****p<0.0001 compared to respective no virus). **E**. Negative correlation between COVID-19 plasma RBD-IgG levels and the RBD-host cell binding. Adjusted R square 0.623 with P=3.58E-07. **F**. Mathematical models used to speculate the potential contribution of ACE2 exosomes (y), residual RBD-IgG (x) and other unknown factors (a, b, c, d,…) to SARS-CoV-2 neutralization.

**Supplementary Fig. S4:**
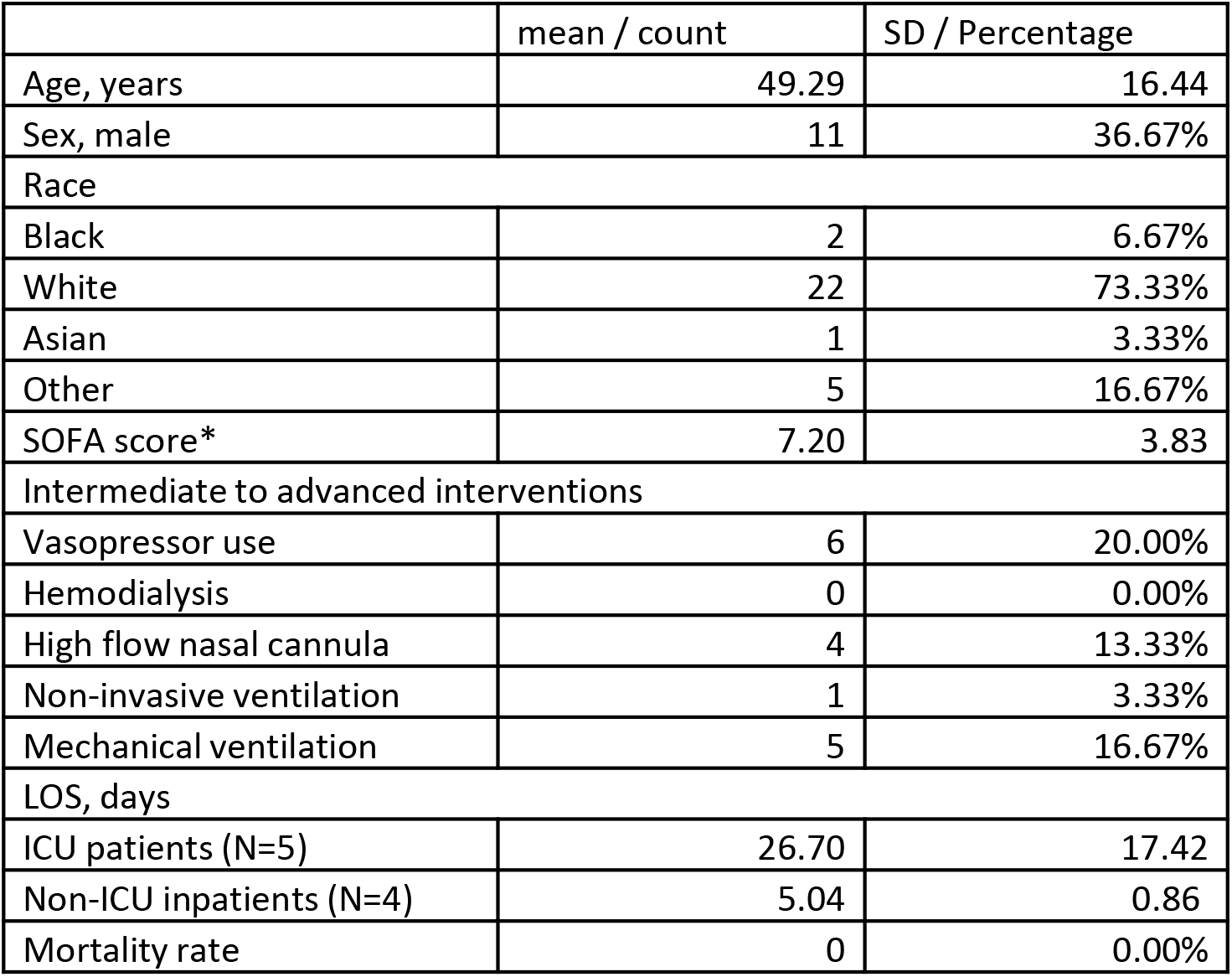
COVID-19 patient table. Summary of convalescent COVID-19 patients from which the plasma samples ACE2^+^ exosomes and RBD-IgG levels were measured (N=30). * The SOFA score was calculated on ICU patients only (N=5). All patients are alive at the time of preparing this manuscript.

